# Microscopy-based chromosome conformation capture enables simultaneous visualization of genome organization and transcription in intact organisms

**DOI:** 10.1101/434266

**Authors:** Andrés M. Cardozo Gizzi, Diego I. Cattoni, Jean-Bernard Fiche, Sergio M. Espinola, Julian Gurgo, Olivier Messina, Christophe Houbron, Yuki Ogiyama, Giorgio L. Papadopoulos, Giacomo Cavalli, Mounia Lagha, Marcelo Nollmann

**Affiliations:** Centre de Biochimie Structurale, CNRS UMR 5048, INSERM U1054, Université de Montpellier, 60 rue de Navacelles, 34090, Montpellier, France; Institut de Génétique Humaine, CNRS UMR 9002, Université de Montpellier, 141 rue de la Cardonille, 34396, Montpellier, France; Institut de Génétique Moléculaire de Montpellier, CNRS UMR 5535, Université de Montpellier, 1919 Route de Mende, 34293, Montpellier, France

**Keywords:** fluorescence microscopy, fluorescent in situ hybridization, genome architecture, topologically associating domains, transcription, Drosophila development, genome organization, chromosome conformation, 4D nucleome, chromatin

## Abstract

Eukaryotic chromosomes are organized in multiple scales, from nucleosomes to chromosome territories. Recently, genome-wide methods identified an intermediate level of chromosome organization, topologically associating domains (TADs), that play key roles in transcriptional regulation. However, these methods cannot directly examine the interplay between transcriptional activation and chromosome architecture while maintaining spatial information. Here, we present a multiplexed, sequential imaging approach (Hi-M) that permits the simultaneous detection of chromosome organization and transcription in single nuclei. This allowed us to unveil the changes in 3D chromatin organization occurring upon transcriptional activation and homologous chromosome un-pairing during the awakening of the zygotic genome in intact *Drosophila* embryos. Excitingly, the ability of Hi-M to explore the multi-scale chromosome architecture with spatial resolution at different stages of development or during the cell cycle will be key to understand the mechanisms and consequences of the 4D organization of the genome.

## Introduction

The study of chromosome organization and transcriptional regulation has been recently revolutionized by the advent of genome-wide sequencing methods. In particular, chromosome conformation capture technologies, such as Hi-C, have revealed that eukaryotic chromosomes are organized into topological-associating domains (Dixon et al., 2012; Nora et al., 2012; Sexton et al., 2012) that can interact together to form active or repressed compartments (Dixon et al., 2015; Lieberman-Aiden et al., 2009). Importantly, disruption of TAD architecture leads to developmental pathologies and disease due to improper gene regulation (Dixon et al., 2016; Hnisz et al., 2016; Spielmann et al., 2018). Thus, determining the role of chromatin architecture in gene regulation has become a key issue in the fields of chromatin biology and transcription.

Recently, single-cell Hi-C and imaging studies showed that chromosomal contacts within and between TADs are highly stochastic and occur at surprisingly low frequencies (Cattoni et al., 2017; Flyamer et al., 2017; Nagano et al., 2017; Stevens et al., 2017). The origin of these heterogeneities has been unclear and may arise from multiple sources, such as variations in transcriptional and/or epigenetic state between cells within multicellular organisms or in cell cultures. Up until now, it has been difficult to detect the origin of these variations, as methods that are able to detect transcriptional output and chromosome organization simultaneously at the single cell level and in the context of an organism have been lacking. In part, this is due to the loss of spatial information in sequencing-based methods.

The study of chromatin architecture and organization by microscopy is limited by several factors. In conventional fluorescence *in situ* hybridization (FISH), the number of spectrally-distinguishable fluorophores in standard microscopes limits the maximum number of genomic loci that can be simultaneously imaged (usually <4). In addition, FISH probes are typically large (>10kb) and difficult to construct, at least in large numbers. These limitations impose a lower limit on the genomic coverage achievable by FISH. Newly developed, low-cost, high-efficiency on-chip DNA synthesis for high coverage Oligopaint FISH have been used to directly label and image entire TADs (Beliveau et al., 2015; Boettiger et al., 2016; Szabo et al., 2018) or TAD borders (Cattoni et al., 2017). These technologies have been recently combined with multiplexed sequential optical imaging to detect hundreds of individual RNA species while still maintaining spatial information (Chen et al., 2015; Eng et al., 2017; Moffitt et al., 2016; Shah et al., 2018), as well as to visualize the segregation of TADs into chromosome territories (Wang et al., 2016). These methods, however, lacked the genomic resolution to detect TADs and to correlate them with transcriptional activity in the context of intact tissues or organisms.

## Design

To overcome these limitations, we introduce a high-throughput, high-resolution, high-coverage microscopy-based technology (Hi-M), capable of simultaneously detecting RNA expression and 3D chromatin organization at the single-cell level with nanometer and kilobase resolutions in intact *Drosophila melanogaster* embryos.

Hi-M microscopy relies on the sequential labeling and imaging of multiple DNA loci in genomic regions spanning hundreds of kilobases (kb). DNA was labeled using oligopaint technologies (Beliveau et al., 2012, 2015). A primary library containing thousands of oligonucleotides targeting multiple genomic loci was designed and produced by high-throughput DNA synthesis (Methods). The subset of oligonucleotides targeting each genomic locus (hereafter barcode) contained unique tails with specific sequences that could be independently read by complementary, fluorescently-labeled oligonucleotide probes (hereafter readout probes; Figures 1A, S1A). An additional barcode (hereafter fiducial barcode), visible in all rounds of hybridization, was used for image registration and drift correction (Methods).

**Fig 1.**
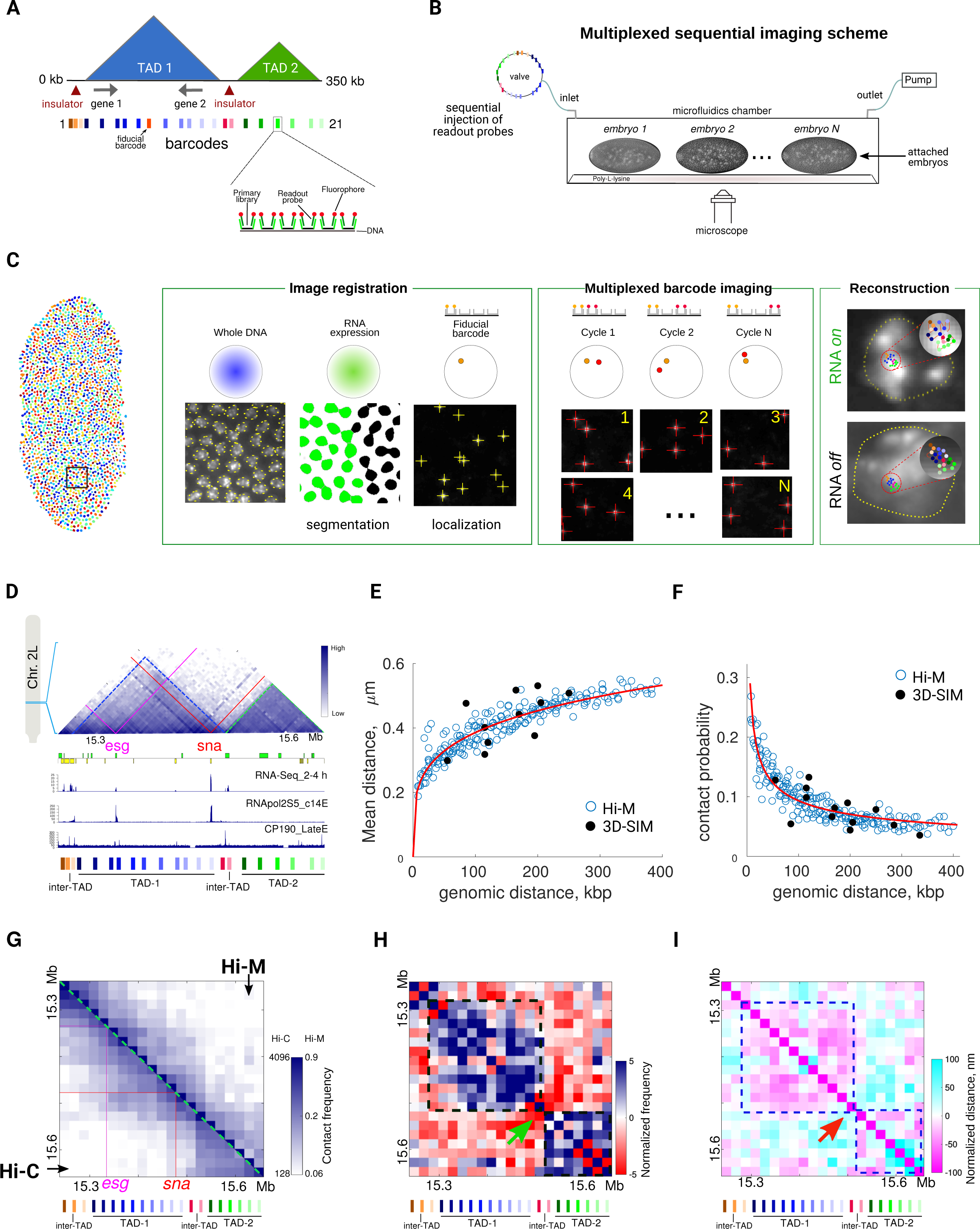
Hi-M enables exploration of chromosome structure in single cells within intact organisms with high genomic coverage and high spatial resolution. A. Schematic representations of an example locus and the labeling strategy are shown. TADs are depicted as blue/green triangles, genes as arrows, insulators as red triangles. Barcodes are indicated as rectangles with a color code following their genomic location. Inset: each barcode is composed of multiple short oligonucleotides with a genome homology region (black), a region for secondary oligo binding (light green) and a fluorescent readout probe (dark green and red). B. Schematic representation of the Hi-M setup. A robotic multi-color, 3D widefield imaging system was coupled to an automatic fluid-handling device (see main text and Methods). Multiplexed barcode imaging is achieved by injecting readout probes sequentially (depicted as colored rectangles). Within the microfluidics chamber, *Drosophila* embryos are attached to a poly-lysine treated coverslip. C. Acquisition and analysis Hi-M imaging pipeline (see Figures S1A-C for more details). An entire embryo is shown where each nuclei is represented by a different color (left). Image registration phase: each nuclei within a field-of-view is segmented based on DAPI staining (left panel). Next, masks were classified according to RNA expression levels as expressing (green) or not expressing (black, middle panel). Fiducial barcodes are imaged, segmented and localized (right panel). Multiplexed barcode imaging phase: readout probes are sequentially injected, imaged, segmented and localized. Reconstruction phase: after drift correction, the 3D position of all barcodes were retrieved for each nuclei in the embryo. Scale bars are indicated in the Figure. D. Hi-C map of nc 14 Drosophila embryos for the region 2L:15.250-15.650 Mb indicating normalized contact probability with a scale going from white (low) to blue (high)(Ogiyama et al., 2018). TADs detected by Hi-C (TAD-1 and TAD-2) are delineated with blue and green dotted lines, respectively. *esg* and *sna* are indicated with magenta and red lines, respectively. Lower panels: genes, RNA-Seq from embryos 2-4 hours of development, Chip-seq from Phospho RNA pol II (S5) from nc 14 embryos and Chip-Seq against CP190. Genomic positions of barcodes are represented as in panel A. E-F. Mean physical distance (E) and absolute contact probability (F) *vs*. genomic distance from Hi-M (blue circles). Black filled circles are data points from our previous study, where we performed pairwise measurements in *Drosophila* embryo cells in a different locus using 3D-structured illumination microscopy (Cattoni et al., 2017). Solid red lines depict a power-law fit with the scaling exponents β = 0.23 ± 0.02 for panel E and α = −0.4 ± 0.02 for panel F. N = 6643. G. Hi-M map and interpolated Hi-C matrix from nc 14 wild-type embryos spanning the ∼400 kb region encompassing *sna* and *esg*. Barcodes are represented as in panel A. *esg* and *sna* are indicated with magenta and red lines, respectively. Features observed in Hi-C at the interpolated resolution and at 5 kb resolution are equivalent (Figure S1J). Relative (Hi-C) and absolute (Hi-M) contact frequencies are color-coded according to scale bar. We estimate the absolute contact probability, as previously described (Cattoni et al., 2017), by integrating the area of the pairwise distance distribution below 120 nm. This threshold value was chosen from the integration of the pairwise distance distribution of a doubly-labeled locus. A good correlation between Hi-M replicates was found (Pearson correlation = 0.92, Figure S1J). N = 6643. H. Normalized Hi-M contact maps from nc 14 wild-type *Drosophila* embryos. Normalization is achieved by dividing observed *vs* expected contact frequencies for equivalent genomic distance. The expected contact frequency is obtained from fit in panel F. Color scale on the right indicates fold enrichment in log scale (positive, blue; negative, red; white, equal). Green arrows indicate chromosomal regions with contact frequencies lower than expected. TADs detected by Hi-C are shown with dotted black lines. Barcodes are represented as in panel A. N = 6643. I. Normalized mean physical distance map from nc 14 wild-type *Drosophila* embryos. Normalization is achieved by subtracting observed and expected distances. The expected distance was obtained from the fit in panel E. Color scale indicates distances lower (magenta) or higher (cyan) than expected. Normalised distances are in nm. Red arrow indicate regions displaying distances higher than expected. TADs detected by Hi-C are shown with dotted black lines. Barcodes are represented as in panel A. N = 6643. See also Figure S1.

Barcodes were imaged using a robotic, fully-automated, four-color microscope coupled to an automated microfluidics system (Figure S1B). The imaging registration phase involved the acquisition of four channel, 3D images of each entire embryo in the chamber to identify embryonic morphology (bright field), nuclei by DAPI staining (blue channel), RNA expression patterns (green channel) and detect the positions of fiducial barcodes (yellow channel, Methods) (Figures 1C and S1C). Next, we performed multiple sequential cycles of hybridization, washing, and imaging of each barcode using a dedicated fluid handling system (Figures 1B-C, Methods). For each hybridization cycle, we performed 3D, two-color imaging of readout probes and fiducial barcodes (Figures 1C and S1A-C and Methods). Sample drift during acquisition was corrected using the fiducial barcode (Figures S1D-E and Methods). A custom-made analysis pipeline was developed for semi-automated image processing and analysis (Methods).

## Results

### Hi-M enables visualization of chromosome structure with high coverage and high resolution

We applied Hi-M to study chromosome organization during early Drosophila development at the time of zygotic genome activation (ZGA). To this purpose, we initially focused on a genomic locus containing the snail (*sna*) and escargot (*esg*) genes, which are among the ∼100 genes expressed during the first wave of ZGA as early as stage 4 (nuclear cycles, nc, 10-13) (Chen et al., 2013). *Sna* and *esg* genes encode zinc finger transcription factors, essential for a variety of processes such as gastrulation, neuroblast specification or stem cell maintenance (Ashraf et al., 1999; Korzelius et al., 2014). In nc 14, the genomic locus encompassing these genes folds into a well-defined TAD (Ogiyama et al., 2018) demarcated by two borders containing class I insulators (CTCF, Beaf-32 and CP190)(Nègre et al., 2010) and housekeeping genes (Figure 1D). To study the 3D organization of this locus by Hi-M, we designed an oligopaint library covering the whole locus with an average distance between barcodes of 17 kb (Figure 1D). This primary library consisted of 2,000 different 142-base pairs oligonucleotides targeting 22 loci around *sna* (21 barcodes and a fiducial barcode).

We performed Hi-M experiments on the *sna-esg* locus in *Drosophila* embryos in nc 14. Dozens of embryos were attached to a poly-L-lysine-coated coverslip and mounted into a microfluidics chamber (Methods, Figure 1B). The overall labeling efficiency was over 60% and did not vary considerably between hybridization cycles (Figure S1G). To validate the method, we first measured the mean physical distances and the absolute probability of interaction between any two loci as a function of their genomic distance (Figures 1E-F, and Methods). These data overlay well with our recent pairwise distance and absolute contact frequency measurements using super-resolution microscopies (Cattoni, et al 2017). Furthermore, there is a good correspondence between replicates, as reflected by similar chromatin polymer properties, contact probabilities, and pairwise distance distributions (Figures S1H-J). It is worth noting that a Hi-M dataset provides a much larger coverage for both types of measurements while being much less time consuming (2 days versus ∼1 year for the datasets displayed in Figures 1E-F).

Next, we constructed a contact probability map from our Hi-M dataset and compared it with published Hi-C data interpolated at the barcodes positions (Figures 1D, S1K and Methods) from equivalent Drosophila embryonic stages (Ogiyama et al., 2018). Hi-C and Hi-M maps are remarkably similar (Fig. 1G). Contact frequencies display a high correlation across several orders of magnitude (Fig. S1L, Pearson correlation coefficient = 0.91). This high correlation between results from two different methods provide a cross validation for both technologies at the length scales probed in this study. Two clearly distinguishable TADs are visible in this locus for both matrices (TAD-1, TAD-2, Figure 1G). TAD-1 contains the *sna* and *esg* genes, and is separated from TAD-2 by a barrier containing highly-expressed housekeeping genes, RNA pol II, class I insulator proteins, and Zelda (Figures 1D, S4D).

To further characterize chromatin organization, we built normalized contact and distance maps implementing a normalization method similar to that performed for Hi-C data (Lieberman-Aiden et al. 2009). Here, we normalized contact frequency and mean spatial distance maps to the expected contact frequencies and spatial distances at each genomic distance as predicted by the power-law scaling fits in Figures 1E-F. The normalized contact map showed enriched interactions within each of the two TADs detected by genomic methods (Figure 1H, dashed boxes) and depleted interactions at TAD borders (Figure 1H, green arrow). Interestingly, pairwise distances within TADs were smaller than expected whereas pairwise distances of loci at TAD borders were higher. Thus, the chromatin fiber appears to be condensed within TADs and decondensed at TAD borders (Figure 1I, dashed boxes and red arrows, respectively).

### Chromosome structure in paired and unpaired chromosomes

*Drosophila* chromosomes display a high degree of homologous pairing in somatic cells (Fung et al., 1998; Joyce et al., 2012). Thus, an important unanswered question is whether chromosome architecture is influenced by pairing. This question cannot be answered by genome-wide methods as they are unable to distinguish between inter- and intra-chromosomal contacts. We took advantage of the capability of Hi-M to discern paired and unpaired homologous chromosomes to compare chromatin structure in each configuration. From the barcodes coordinates in each nucleus, we classified chromosomes as being ‘paired’ if all barcodes in that nucleus are detected not more than once. Instead, a nucleus contains ‘unpaired’ chromosomes when at least one barcode is detected more than once in that nucleus. In this definition, partially or completely unpaired chromosomes are classified as ‘unpaired’ (Figure S2A). In our Hi-M data, the frequency of barcode pairing was ∼70 % for nc 14 nuclei (Figure S2B), in good agreement with previously published results (Bateman and Wu, 2008; Fung et al., 1998). Normalized distance maps and contact matrices for paired and unpaired chromosomes were qualitatively very similar (Figures 2A and S2C-D). To quantitatively compare these two configurations, we performed a multi-scale correlation analysis of Hi-M distance maps (Methods). This analysis revealed that Hi-M matrices are almost identical (correlation = 1) at low resolution and retained a large, although reduced, similarity even at the highest resolution (correlation = 0.6, Figure 2B). This indicates that the overall organization of chromatin into TADs is partially similar in paired and unpaired chromosomes (see discussion in Figure S2E). Consistently with this picture, mean pairwise distances between paired and unpaired chromosomes were highly correlated (Figure 2C). Remarkably, however, pairwise distances in unpaired chromosomes were in most cases larger than the corresponding distance for paired chromosomes (Figure 2C). These results suggest that chromatin folding into TADs is similar between paired and unpaired chromosomes but there is certainly an overall compaction of chromatin upon pairing or a decompaction upon un-pairing.

**Fig 2.**
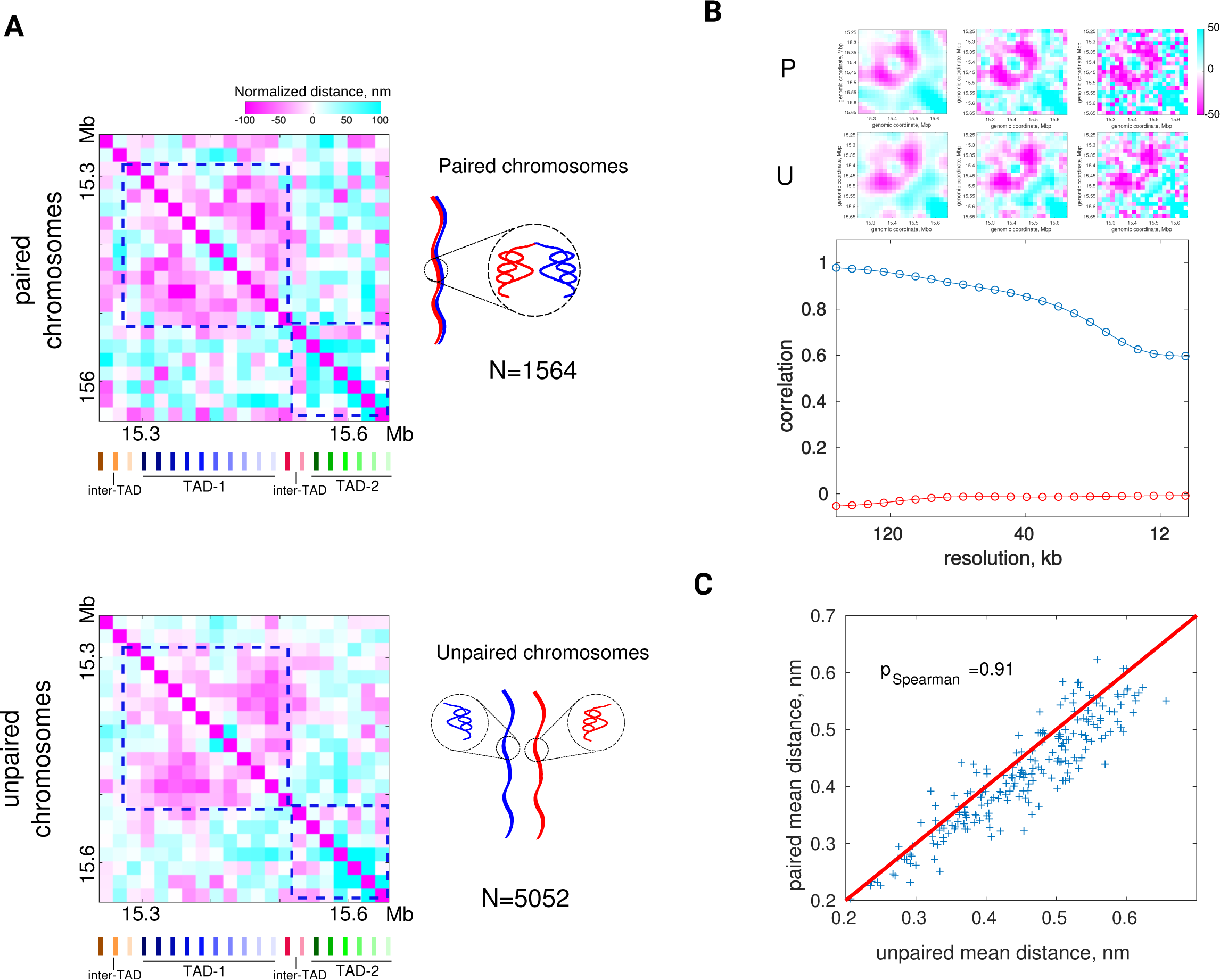
Chromosome architecture in unpaired and paired homologous chromosomes. A. Hi-M normalized distance maps from paired (top panel) or unpaired (lower panel) homologous chromosomes. Barcodes are represented as in Fig 1A. Schematic representation indicate paired/un-paired homologous chromosomes (top-right and bottom-right panels). Number of examined nuclei, N= 1564 (paired), N=5052 (unpaired). B. Top panels, Hi-M interpolated normalized distance maps at three different resolutions as examples for paired (P) and unpaired (U) chromosomes. Scale bar is on the right. Bottom panel, multi-scale correlation of Hi-M normalized distance maps for paired and unpaired chromosomes. Blue circles represent the correlation of maps between paired and unpaired chromosomes at different resolutions. Red circles represent the correlation between randomized matrices as a function of resolution. N as in panel A. C. Paired chromosomes mean pairwise distances vs. unpaired chromosome mean pairwise distances, represented as blue crosses. The red line represents a slope equal to 1. Note the tendency of unpaired chromosomes pairwise distances to be higher than the corresponding for paired chromosomes. Pearson correlation coefficient = 0.91. N as in panel A. See also Figure S2.

### Chromosome organization changes during cell-cycle and development

Recent genome-wide studies showed that chromosome organization into TADs changes during the cell cycle (Hug et al., 2017; Nagano et al., 2017; Naumova et al., 2013). To study whether we could observe such changes by microscopy, we performed Hi-M in embryos undergoing mitosis (Figure 3A). We observed that TADs were no longer discernible at this phase of the cell cycle and that the frequency of genomic contacts was almost independent of genomic distance at short scales (<400 kb, Figures 3B-D, S3D), reflecting a lack of hierarchical organization during mitosis. Overall, these results are in excellent agreement with published Hi-C data (Hug et al., 2017; Nagano et al., 2017; Naumova et al., 2013) (Figures 3B, S3A). Normalized distance maps indicate a low correlation between physical and genomic distances (Figure 3D), consistent with irregular, intermingled loops forming a uniform-density, phase-like structure as previously proposed (Naumova et al., 2013; Nishino et al., 2012).

**Fig 3.**
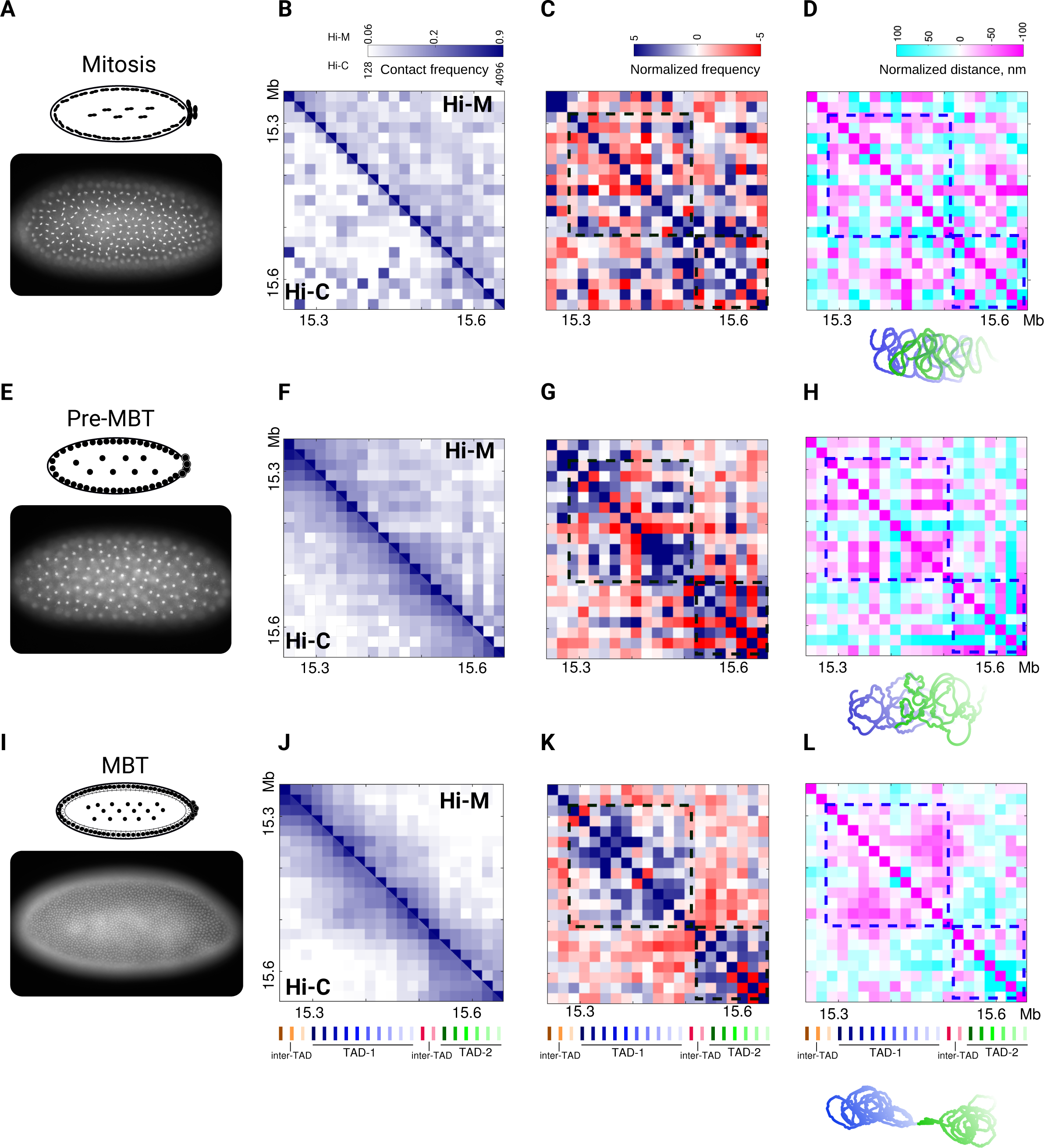
In situ, single-cell exploration of chromosome conformation during the cell cycle and development. A,E and I. Top, schematic representation of an embryo undergoing mitosis (A), nc 9-13 (pre-MBT) (E) and nc 14 (MBT) (I). Bottom, DAPI image of a representative embryo of the corresponding stage. Scale bar = 50 μm. B,F and J. Hi-M map (upper right triangle) and interpolated Hi-C matrix (lower left triangle) from embryos undergoing mitosis (B), nc 9-13 (pre-MBT) (F) and nc 14 (MBT) (J). Barcodes are represented as in Figure 1A. Features observed in Hi-C at the interpolated resolution and at 5 kb resolution are equivalent (Figure S3A). Relative (Hi-C) and absolute (Hi-M) contact frequencies are color-coded according to scale bar. Number of examined nuclei, N= 1430 (mitosis), N=2933 (nc 12-13), N=6643 (nc 14). C,G and K. Normalized Hi-M contact maps from embryos undergoing mitosis (C), nc 9-13 (pre-MBT) (G) and nc 14 (MBT) (K). Color scale on top as in Figure 1H. TADs assigned by Hi-C are shown with dotted black lines. Barcodes are represented as in Figure 1A. N as in previous panel. D,H and L. Normalized mean physical distance maps from embryos undergoing mitosis (D), nc 9-13 (pre-MBT) (H) and nc 14 (MBT) (L). Color scale on top as in Figure 1I. TADs observed in nc 14 are delineated with dotted blue lines. Barcodes are represented as in Figure 1A. Schematic representations for each developmental state are shown below matrices. Blue and green represent the chromatin fiber in each of the two TADs detected by Hi-C. N as in previous panel. See also Figure S3.

Next, we used Hi-M to investigate whether chromosome organization changed during the mid-blastula transition (MBT) at the onset of the major wave of zygotic transcription occurring at nc 14. For this, we performed Hi-M in embryos in nc 12-13 (pre-MBT) and nc 14 (MBT) (Figures 3E, 3I). Hi-M maps displayed a good correspondence with Hi-C maps (Figures 3F-G, 3J-K). Notably, TADs emerged at the onset of the ZGA (Figures 3G, 3K), consistent with previous Hi-C studies (Figures 3F, 3J, and S3B-C) (Hug et al., 2017; Ogiyama et al., 2018). Hi-M contact frequencies decayed more dramatically with genomic distance in nc 14 than in previous nuclear cycles, consistent with a change in the overall organization of the chromatin fiber occurring at this stage (Figures S3D-F). These local changes in chromatin organization were also clearly seen in normalized mean distance maps, where a progressive condensation of chromatin in TADs between nc 12-13 and nc 14 can be observed (Figures 3H and 3L). Next, we calculated the coefficient of variation (ratio between standard deviation and mean), a measure of heterogeneity in the cell population. We observed that maps of coefficient of variation are correlated to emergence of TADs (Figures S3G-I). Interestingly, at nc 14 the coefficient of variation was: ∼1 inside TADs, low (∼0.5) in regions of the map encompassing interactions between TADs, and high (∼1.5) in gene-rich regions (see RNA-seq and RNA pol II profiles in Figure 1D). In contrast, this pattern was disrupted in early embryos (nc 12-13) or in mitotic cells where TAD architecture was absent (Figure S3G-H). Altogether, these results indicate that heterogeneity in chromosome architecture is modulated by structural and functional features of the genome.

### Simultaneous measurement of TAD organization and transcriptional activity in single nuclei

Several studies correlating chromosome organization and transcription have suggested that TAD organization changes upon transcriptional activation (Cruz-Molina et al., 2017; Hug et al., 2017; Phanstiel et al., 2017; Stadhouders et al., 2018). However, due to intrinsic limitations in Hi-C technologies, it has been so far impossible to detect chromosome organization and transcriptional state at the same time in single cells. We set out to test whether Hi-M could be adapted to perform this measurement. For this, we included an RNA labeling step in our imaging pipeline (Methods) that allows us to detect which nuclei in the embryo are transcriptionally active (Figure 4A). Specifically, we labeled and imaged *sna* transcripts in whole embryos, and detected its characteristic ventral expression pattern in nc 14 (Alberga et al., 1991) (Figure 4A).

**Fig. 4.**
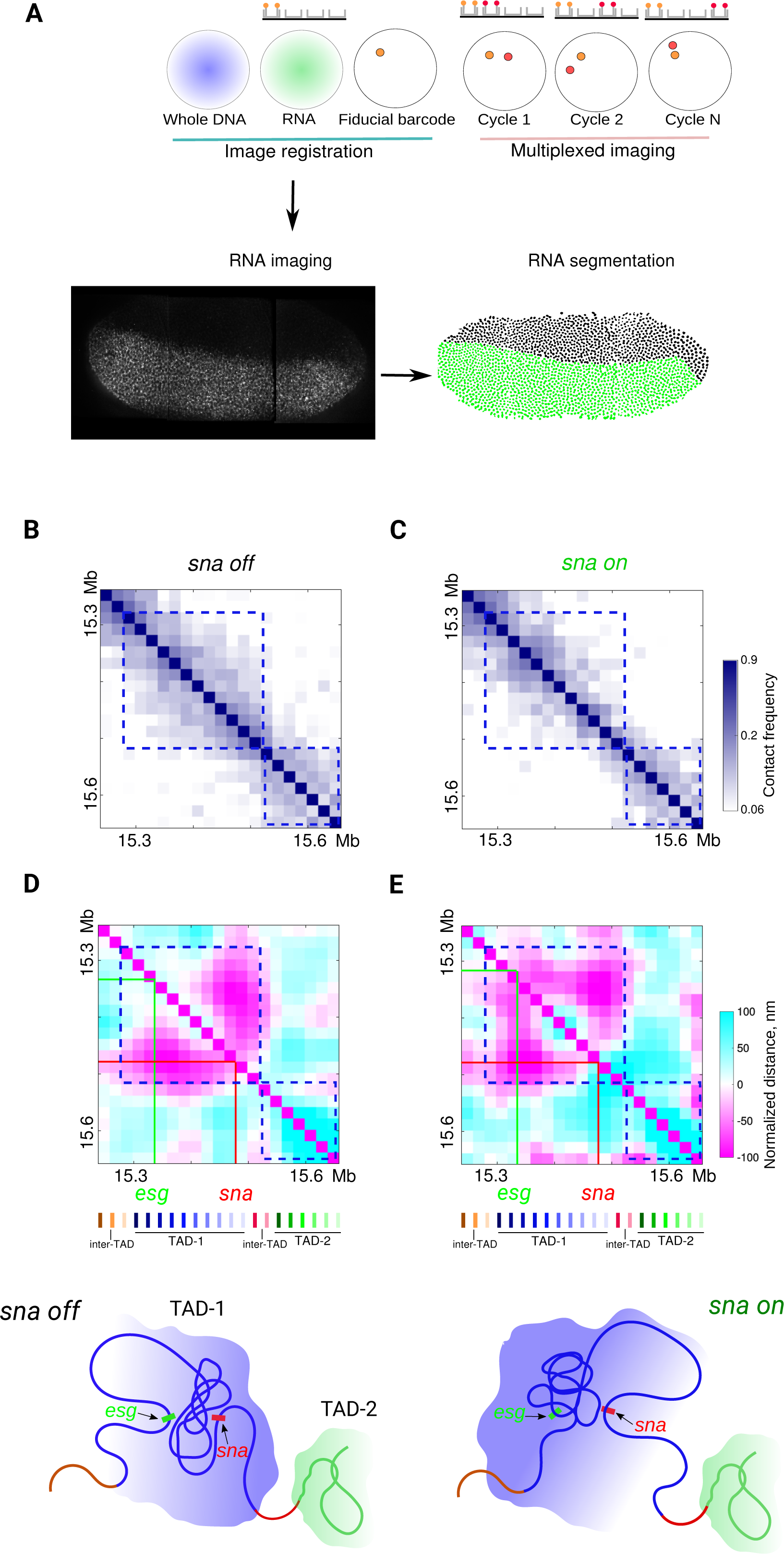
Simultaneous detection of chromosome organization and transcription by Hi-M. A. Single-cell RNA expression detection. Top: scheme depicting the Hi-M experimental design. Bottom left: Representative full embryo *sna* RNA image obtained from the image registration step. Image is composed of a mosaic of three fields of view. Scale bar= 20 μm. Bottom right: result of nuclei segmentation and RNA state assignation for the full embryo. Nuclei expressing *sna* are in green whereas nuclei not expressing *sna* appear in black. For the displayed embryo in the example, 63% of nuclei are expressing sna, although this percentage depends on the orientation. B-C. Hi-M absolute contact matrices for nuclei expressing (C) or not (B) expressing *sna*. Absolute contact frequency is color-coded according to scale bar as goes from 0 to 0.1. Barcodes are represented as in Figure 1D. N=4402 (off nuclei examined) and N=2265 (on nuclei examined). D-E. Normalized Hi-M distance maps for nuclei expressing (E) or not (D) expressing *sna*. Color scale indicates distances shorter (magenta) or higher (cyan) than expected (scale in nm). Solid lines represent the positions of *sna* and *esg*. Barcodes are represented as colored as in Figure 1A. On the bottom, schematic representations of TAD organization in transcriptionally *on/off* nuclei are shown. TAD-1 is represented in blue and TAD-2 in green. N as in previous panel.

Hi-C maps at the *sna* locus contains two TADs (TAD-1 and TAD-2), with TAD-1 displaying looping interactions between regions surrounding *sna* and *esg* (Figure S4A). Surprisingly, TAD-1 could be clearly discerned in nuclei where *sna* was repressed but not in transcriptionally active nuclei (Figures 4B-C). These changes are also clearly observed in normalized contact maps (Figure S4B). This observation strongly suggests that chromatin organization into TADs is disrupted by transcriptional activation.

To gain further insight into the structural changes that may underpin this difference in TAD organization, we calculated the normalized median distance maps for *sna* positive nuclei and for their neighbouring inactive nuclei. The patterns of chromatin folding within the largest TAD (containing *sna*) showed clear changes upon *sna* expression (Figures 4D-E, S4C). Strikingly, chromatin structure in the vicinity of *sna* was locally decondensed in nuclei exhibiting active *sna* transcription. In nc 14, the expression patterns of *sna* and *esg* do not overlap in space (Hemavathy et al., 2004). Thus, the local chromatin decondensation in the vicinity of *esg* in nuclei that are not transcribing *sna* is consistent with active *esg* transcription leading to local chromatin decondensation.

Hi-C maps from nc 14 display specific looping interactions between regions surrounding *esg* and *sna* (Figure S4A). However, Hi-C cannot discern whether these contacts depend on the transcriptional status of these genes. Surprisingly, by Hi-M we observe dramatic changes in the internal organization of TAD-1 upon *sna* activation, with genomic regions around *sna* and *esg* being closer than expected in both active and inactive nuclei (Figures 4D-E, S4B). Exploration of existing Chip-seq datasets showed that the region surrounding and encompassing these genes is occupied by active (GAGA-factor, Zelda, CP190) (Blythe and Wieschaus, 2015) as well as inactive marks (Polycomb group proteins, H3K27me3) (Figure S4D). Thus, we hypothesize that different networks of contacts are responsible for the distinct patterns of interactions visualized by Hi-M in active and inactive nuclei.

## Discussion

We developed a new method — Hi-M — based on sequential, multiplexed, high-throughput hybridization and imaging of oligopaint probes to visualize chromosome organization and transcriptional activity simultaneously in single cells while preserving tissue context. We validated our method by comparing our results with existing microscopy and Hi-C datasets, and by showing that TADs detected by Hi-M are disrupted during mitosis and emerge at the onset of the ZGA during embryonic development. Overall, these experiments strongly support the ability of Hi-M to capture chromosome conformations at the single cell level in a variety of experimental conditions. In addition, we used Hi-M to directly show that chromosome pairing leads to a general compaction of chromatin without an overall change in TAD organization; and that distance distributions within TADs dramatically change upon transcriptional activation.

The degree of pairing between homologous chromosomes in *Drosophila* has been previously characterized (Bateman and Wu, 2008; Cattoni et al., 2017; Fung et al., 1998; Joyce et al., 2012), however the organization of chromatin within paired or unpaired homologous chromosomes remained inaccessible to conventional microscopy imaging. Hi-M revealed that paired and unpaired chromosomes share equivalent TAD architectural features but that the latter have a less compact folding. Interestingly, during the revision process of this work, a new Hi-C study confirmed our Hi-M predictions for early *Drosophila* embryos showing the conservation of domains in paired and unpaired chromatin (Erceg et al., 2018).

Upon mitosis the decrease of insulation at borders leads to loss of chromatin compartilization in metazoans (Nagano et al., 2017; Naumova et al., 2013). Hi-M retrieved equivalent results by directly evaluating the absolute frequency of interaction at the single-nuclei level. Moreover, the homogeneous variance distribution of distances (Figure S3G) during mitosis obtained by Hi-M, is compatible with the recently proposed model of helical arrangements and nested loops of mitotic chromosomes (Gibcus et al., 2018). Hi-M opens many new possibilities, such as the determination of 3D chromatin architecture at different mitotic stages (prophase, metaphase, anaphase) to decipher at which of these stages TAD organization is regained and the mechanisms involved in this process.

Early evidence suggesting large-scale chromatin structure changes upon transcriptional activation (Tumbar et al., 1999) has been recently confirmed by genome wide approaches focusing on genome activation during development (Du et al., 2017; Hug et al., 2017; Ke et al., 2017). Hi-M reproduced satisfactorily previous genome wide findings of chromosome organization during the zygotic awakening of *Drosophila* and gave additional insight into the organization of chromatin. Particularly, we show that not only changes in the frequency of interaction between loci accompanies gene activation as it has been previously suggested (Hug et al., 2017), but also a full reshaping of the 3D organization of the TAD, compatible with recently proposed models based on kinetic measurements of transcription and local reorganization of chromatin (Chen et al., 2018)(Shah et al., 2018).

Excitingly, Hi-M can be widely applied to detecting single-cell chromosome organization and transcription in cultured cells, from bacteria to mammals, or in multicellular organisms and tissues. We note that a key advantage of our method is that it would also allow for the combination of single nuclei analysis with spatial mapping of the relationship between structure and function, i.e. chromatin architecture and gene expression, in fly embryos and in more complex tissues. Thus, this novel technology has the potential to revolutionize the study of chromosome architecture in many fields and at many different scales.

## Limitations

Here, we show that Hi-M can simultaneously label ∼21 distinct DNA loci. Current developments in liquid handling technologies are being made to increase this number by about one order of magnitude, which would allow for an overall increase in the size of the region that can be probed by Hi-M in similar time-scales. Further optimization of Hi-M will involve simultaneous acquisition of several colors which would shorten considerably the acquisition time by a factor of 2-3 (Moffitt et al., 2016). Use of combinatorial labelling schemes should make it possible to considerably increase the number of detected DNA loci without increasing the number of hybridization cycles (Chen et al., 2015; Shah et al., 2018).

A second current limitation of Hi-M is that it can detect a single RNA species, as it requires the use of digoxigenin-labeled RNA probes. We envision that it should be relatively straightforward to increase the number of RNA species by using orthogonal chemistries (e.g. biotin). This approach, however, will be limited to a few RNA species. To considerably increase the number of detected RNA species will likely require that RNA species are detected prior to DNA detection using adaptations of multiplexed RNA detection protocols (Chen et al., 2015).

A third limitation of Hi-M is the size of probes (currently ∼4 kb, 70 primary oligos). We envision that the use of fewer primary oligos (∼20-30) should ensure specificity of detection and reduce the size of the probe to ∼1.5-2kb. Further increases in the number of fluorophores per primary oligonucleotide could enable sub-kb resolutions. It is, however, important to bear in mind that a reduction in probesize will have to be balanced by a reduction in the genomic frequency of probes or by a reduction in the size of the genomic region being probed. The minimal inter-probe distance used in this study was ∼10 kb. We note that further reducing the inter-probe distance may require further improvements in the localization precision and drift correction. Finally, our current implementation of Hi-M uses widefield microscopy. This method is ideally suited for detection in early Drosophila embryos where cells are organized in a 2D layer, but not for samples with more complex architectures (such as late stage embryos). For these architectures, a confocal imaging scheme would be better suited, and will be explored in future.

## Acknowledgments

The authors thank Carola Fernandez for providing the *sna* probe and for the help with RNA labeling experiments. This project has received funding from the European Research Council (ERC) under the European Union’s Horizon 2020 research and innovation programme (Grant Agreement No 724429). This work also benefited from support by the Labex EpiGenMed, an «Investments for the future» program, reference ANR-10-LABX-12-01. We acknowledge the France-BioImaging infrastructure supported by the French National Research Agency (ANR-10-INBS-04, «Investments for the future»). This work was also supported by the ERC SyncDev starting grant to M.L.

## Authors contribution

A.M.C.G., M.L. and M.N. designed experiments. A.M.C.G., D.I.C., S.E., J.G. and O.M. conducted research. J.B.F. designed and built microscopy setup and acquisition software. M.N. developed software for image analysis. A.M.C.G., J.G., and M.N. analyzed data. A.M.C.G. and M.N. designed oligopaint probes. J.G and O.M. performed fly handling. C.H. synthesized and purified oligopaints libraries. Y.O., G-L. P. and G.C. performed Hi-C experiments and analysis. A.M.C.G., D.I.C, and M.N. wrote the manuscript. All the authors reviewed and commented on the data.

## Declaration of interests

The authors declare no competing financial interests.

See also Figure S4.

## STAR Methods

### CONTACT FOR REAGENT AND RESOURCE SHARING

Further information and requests for resources and reagents should be directed to and will be fulfilled by the Lead Contact, Marcelo Nollmann.

### EXPERIMENTAL MODEL AND SUBJECT DETAILS

Oregon-R w^1118^fly stocks were maintained at room temperature with natural light/dark cycle and raised in standard cornmeal yeast medium. Following a pre-laying period of 16-18 hours in cages with yeasted 0.4% acetic acid agar plates, agar plates were changed for new ones so flies can lay eggs during 1.5 hours on the new plates. Embryos were then incubated at 25 °C for an extra hour with 2.5 h of total developmental time at the time of fixation.

## METHOD DETAILS

### Drosophila embryo collection

Embryos collection were as described (Trcek et al., 2017). Briefly, embryos were dechorionated with bleach for 5 min and thoroughly rinsed with water. They were fixed in fixation buffer (1:1 mixture of 4% methanol-free formaldehyde in PBS and Heptane) by agitating vigorously for 15 sec and then letting stand the vial for 25 min at RT. The bottom formaldehyde layer was replaced by 5 ml methanol and embryos were vortexed for 30 s. Embryos that sank to the bottom of the tube, devitellinized, were rinsed three times with methanol. Embryos were stored in methanol at −20 °C until further use.

### Oligopaint libraries

Oligopaint libraries were obtained from the Oligopaint public database (http://genetics.med.harvard.edu/oligopaints) and consisted of unique 42-mer sequences with homology to the genome. Probe density was around 10-15 probes per kb. We selected 22 genomic regions of interest (barcodes from now on) in the *sna* locus (2L:15244500..15630000 Drosophila release 6 reference genome), spanning a total of ∼400 kb with an average distance between barcodes of 17 kb. For each barcode, we obtained seventy-five probes, covering between 4-6 kb. See Table S1 for the coordinates of all barcodes used. One of the barcodes was selected as the fiducial barcode for image registration and drift correction.

Each oligo in our template primary probe library contained 5 regions: (from 5’ to 3’) i) a 21-nucleotide (nt) forward universal priming region for library amplification (5’-CGCTCGGTCTCCGTTCGTCTC), ii) a 30/32-nt readout region for binding of a complementary fluorescent secondary probe unique for each barcode (see Table S2 for all of readout region sequences), iii) the 42-nt genome homology region for *in situ* hybridization to the target chromosomal DNA sequence, iv) a duplication of 30/32-nt readout region to allow for the binding of a complementary second secondary probe and v) a 21-nt reverse universal priming region (5’-GCTGAACCCTGTACCTAGCCC). An oligo pool with all the oligonucleotides used in this study (∼2000) was ordered from Twist Bioscience (San Francisco, USA). The 4-step procedure used to amplify the oligopaint probes was as described elsewhere (Wang et al., 2016). Briefly, it consists of a i) limited-cycle PCR, with pairs of PCR primers (BB287-FWD: 5’-CGC TCG GTC TCC GTT CGT CTC/ T7+BB288-REV: 5’-TAA TAC GAC TCA CTA TAG GGT TGG GCT AGG TAC AGG GTT CAG C) targeting the 21-nt forward and reverse priming regions. The reverse primer also contained an additional T7 promoter sequence (5’-TAATACGACTCACTATAG). This allows for ii) further amplification via T7 *in vitro* transcription using T7+BB288-REV primer, which were then iii) converted back to single-stranded DNA oligo probes via reverse transcription using BB287-FWD primer. Finally, iv) the intermediate RNA products were removed with alkaline hydrolysis and DNA oligo probes were purified via column purification.

Secondary readout 32-mer fluorescently-labeled oligonucleotides (fluorescent readout probes, see Table S3) were synthesized by Integrated DNA Technologies (IDT; Coralville, USA). We employed 22 unique-sequence oligos, 21 of which have a cleavable Alexa-647 attached to the oligo whereas the one used for fiducial barcodes had a non-cleavable Rhodamine fluorophore. The cleavable bond was a disulfide linkage that was removable with the mild reducing agent Tris(2-carboxyethyl)phosphine (TCEP), and allowed us to eliminate the fluorescence of a particular barcode from one cycle to the next (Moffitt et al., 2016). The whole set of Oligopaints used are in Table S4.

### RNA-FISH probe preparation

Sna probe was previously used in (Lagha et al., 2013). The full length sna gene 1.6 kb (Dmel_CG3956, 15476621..15478176 Drosophila release 6 reference genome, flanking sequences 5’-ATTTAATTCTTCTCTTTAAGC-3’ / 5’-GGGTAAATCGGGAGATCGGCG-3’) was cloned into a pBluescript II SK (+) vector, cut by NotI to linearize the vector and in vitro transcribed using T7 RNA polymerase in the presence of digoxigenin haptenes. RNA probe produced this way was then treated with carbonate buffer at 65 °C for 5 min.

### RNA-FISH coupled with TSA

The *in situ* hybridization protocol for RNA detection was that of (Kosman et al., 2004) with minor modifications. Methanol-stored, fixed embryos were rinsed once with fresh methanol, then passed through 1 ml of 1) 50% methanol, 50% ethanol (once), 2) 100% ethanol (5 times for 3-5 min). Incubations were made in all cases for the indicated times at RT on a rotating wheel for each step unless otherwise specified. After two washes with methanol, embryos were incubated with a 1:1 dilution of methanol with 5% formaldehyde in PBT (PBT=0.1% Tween-20 PBS) for 5 min and then rinsed with 5% formaldehyde PBT to help remove methanol. Next, embryos were post-fixed with 5% formaldehyde in PBT for 25 min. Embryos were rinsed twice with PBT, incubated 4 times with PBT during 15 min and permeabilized 1 h with 0.3% Triton in PBS. After 3 five-min washes with PBT, embryos were incubated for 10 min with a 1:1 dilution of PBT with RHS (RHS= 50% formamide, 2X SSC, 0.1% Tween-20, 0.05 mg/ml heparin, 0.1 mg/ml salmon sperm). Then, embryos were incubated with RHS at 55 °C for 10 min, solution changed and incubated during 45 min and then a final incubation of 1 h 15 min. All incubations at 55 °C were made in a Thermomixer with 900 rpm agitation. Next, 2 µL of digoxigenin-labeled RNA probe was diluted in 250 µL of RHS, denatured by heating at 85 °C for a maximum time of 2.5 min and then placed in ice for at least 5 min. Embryo media at 55 °C was removed and the probe-containing RHS was immediately added directly from the ice. Embryos were kept 16-20 h at 55 °C for RNA hybridization. The second day, RNA probe was removed, embryos were washed 4 times at 55 °C with RHS for 30 min each. After one 10 min wash at RT with a 1:1 dilution of RHS with PBT, 3 incubations with PBT for 20 min were made. Then, a saturation step was performed with 2X blocking solution (10X Blocking solution= 10% (w/V) blocking reagent Sigma #11096176001, 100 mM Maleic acid, 150 mM NaCl, pH=7.5) for 45 min, and then the activity of endogenous peroxidases eliminated by incubating with 1% H_2_O_2_ in PBT for 30 min. Finally, after rinsing twice with PBT, embryos were incubated overnight at 4 °C with sheep anti-digoxigenin conjugated with POD (Sigma-Aldrich cat #11207733910) with 1:500 working dilution in PBT. The next day, embryos were rinsed twice with PBT and then washed 5 times with PBT for 12 min each time. For the tyramide signal amplification (TSA), embryos were incubated 30 min with a dilution of 5 µL of Alexa-488 coupled to tyramide dissolved in DMSO (Stock initially dissolved in 150 µL of DMSO to obtain a 100X solution, Invitrogen cat#B40953) in 500 µL of PBT. Next, H_2_O_2_ was directly added to the tube to a final concentration of 0.012% during another 30 min. Embryos were finally washed 3 times with PBT for 5 min.

### Hybridization of primary oligopaint library

Embryos were resuspended by sequential dilutions of methanol with 0.1% V/V Tween-20 PBS (PBT). RNA labeled embryos were already in PBT, so this step was omitted. Next, embryos were RNAse treated during 2 h, permeabilized 1 h with 0.5% Triton in PBS and rinsed with sequential dilutions (20 min each) of Triton/pHM buffer to 100% pHM (pHM = 2X SSC, NaH_2_PO_4_ 0.1M pH=7, 0.1% Tween-20, 50% formamide (v/v)). Then, 225 pmols of the barcode probes were diluted in 30 µL of FHB (FHB = 50% Formamide, 10% dextran sulfate, 2X SSC, Salmon Sperm DNA 0.5 mg ml^-1^). Barcodes and embryos were preheated at 80 °C during 15 min. The supernatant of pHM buffer was completely removed from embryos and 30 µL of barcodes-containing solution were rapidly added. Mineral oil was added on top of the mix to avoid evaporation and the sealed tube was deposited in a water bath at 80°C. Immediately, the water bath was set to 37 °C and let cooling down overnight. The next day, oil was carefully removed and embryos were washed two times at 37 °C during 20 min with 50 % formamide, 2X SSC, 0.3 % CHAPS. Next, embryos were sequentially washed at 37 °C for 20 min with serial dilutions of formamide/PBT to 100% PBT. An additional (optional) crosslink step with PFA 4% was performed and embryos were washed and resuspended in PBS. For the fiducial barcode readout, hybridization was performed in the bench before mounting the sample into the flow chamber. The sample was incubated with Rhodamine-labeled readout probe in hybridization buffer for 30 min at room temperature. Next, the sample was washed and kept in PBT. An additional fixation step could be performed as described before. Finally, embryos were stained with 0.5 µg ml^-1^ of DAPI for 20 min, washed with PBT and stored at 4 °C until imaging.

### Robotic microscope setup

All experiments were performed on a home-made imaging setup built on a RAMM modular microscope system (Applied Scientific Instrumentation - USA). The RAMM module was equipped with a 60x Plan-Achromat water-immersion objective (NA=1.2, Nikon – Japan). The objective lens was mounted on a closed-loop piezoelectric stage (Nano-F100, Mad City Labs Inc. - USA) allowing for a fine control of the focus and the acquisition of z-stacks when imaging embryos. A two-axis translation stage was used to move the sample laterally and select embryos before each experiment (MS2000, Applied Scientific Instrumentation – USA). Four lasers with excitation wavelengths of 405 nm, 488 nm, 561 nm, and 641 nm were used for fluorescence imaging (OBIS-405/488/640 and Sapphire-LP-561, Coherent – USA). Laser beams were combined by a series of dichroic mirrors (LaserMUX™, Semrock – USA), individually controlled by an acousto-optic tunable filter (AOTFnC-400.650, AAopto-electronic – France) and focused onto the back-focal plane of the objective through one of the excitation ports of the RAMM. Excitation and emission wavelengths were separated using a four-band dichroic mirror (zt405/488/561/640rpc-UF2, Chroma – USA) and the fluorescence signal was further filtered using a four-band filter (zet405/488/561/640m, Chroma – USA) before being imaged on a sCMOS camera (ORCA Flash 4.0V3, Hamamatsu – Japan). The final pixel size was 100 nm. A home-made autofocus system was used to correct for axial drift in real-time and maintain the sample in focus while imaging. This was achieved as follows. Along with a separate path, a 785 nm infrared laser beam (OBIS-785, Coherent – USA) was focused onto the back focal plane of the objective and reached the coverslip-sample interface in total internal reflection conditions. The position of the reflected beam was measured by a position-sensitive detector (OBP-A-4H, Newport – USA) and any variations in the objective-sample distance were corrected through the z-positioning piezo stage using a proportional-integral-differential feedback loop.

For sequential hybridizations, a fluidic system similar to the one described by was designed (Chen et al., 2015). The sample was mounted in a FCS2 flow chamber (Bioptechs – USA). Buffers and probe handling were computer-controlled using a combination of three eight-way valves (HVXM 8-5, Hamilton – USA) and a negative pressure pump (MFCS-EZ, Fluigent – France) (Figure S1B). The flow rate was monitored in real-time using an online flow unit (FLU_L_D, Fluigent – France), allowing for a precise control of injected volumes.

All instruments, including camera, stages, lasers, pump, and valves were controlled using a custom-made software package developed in LabView 2015 (National Instrument – USA). This software controlled and synchronized multi-color 3D imaging and automated fluid handling.

### Sequential image acquisition

Embryos labeled with a primary oligopaint library (see above) were attached to a poly-L-lysine coated coverslip and mounted into the FCS2 flow chamber, connected to the fluidics system and secured to the translation stage (see microscope setup). The fluidics system had 21 tubes connected and distributed as follows: 1 tube with 50 mL of washing buffer (WB, 2×SSC, 40% v/v formamide), 1 tube with 50 mL of 2x SSC, 1 tube with 20 mL of imaging buffer (IM, 1xPBS, 5 % w/v glucose, 0.5 mg/mL glucose oxidase and 0.05 mg/mL catalase), 1 tube with 50 mL of chemical bleaching buffer (CB, 2X SCC, 50 mM TCEP hydrochloride) and 17 tubes with 2.5 mL of each readout probe solution (25 nM readout probe, 2×SSC, 40% v/v formamide).

Embryos remained firmly attached when confronted with constant flow (∼200 uL/min) for more than 72 hs, largely exceeding the total imaging time required for a single experiment (Figure 1SC). Following embryo selection, embryos were segmented into a mosaic of several fields of view (FOV of size 200 × 200 µm). First, bright field images were taken for all FOV. Next, DAPI and RNA staining were imaged together with fiducial barcodes by exciting at 405, 488 and 561 nm respectively. Fiducial barcodes hybridized to a Rhodamine-labeled readout probe before mounting the sample. Z-stacks of 15 µm with steps of 250 nm were acquired for all channels. Then, the robotic microscope controlled the sequential hybridization and imaging procedure (see Figure S1B-C). The flow chamber was initially flushed with 1.7 mL readout hybridization mixture over the span of 15 min to exchange buffers fully and ensure to saturate binding of readout probes. Next, the sample was washed with 2 mL of wash buffer for 18 min. Then 1.5 mL of 2X SSC were flushed during 15 min and finally 0.9 mL of imaging buffer was injected in 5 min. At this stage, flow was stopped, and ∼100 FOVs were imaged in two channels by exciting at 561 and 641 nm to image fiducial barcodes and readout probes, respectively (see Figure S1C). After imaging, the fluorescence of the readout probes was extinguished using chemical bleaching by flowing 2 mL of CB buffer for 15 min. The Rhodamine-labeled fiducial barcode was insensitive to chemical removal. In all cases, the flow speed varied between 0.1 and 0.25 mL/min. After chemical bleaching, the chamber was flushed with 2 mL of 2× SSC for 5 min and a new hybridization cycle started. A standard experiment required between 30 and 40 h depending on the number of probes and number of imaged FOV.

All buffers were freshly prepared and filtered for each experiment. To avoid degradation by oxygen, the imaging buffer used for a single experiment was stored under a layer of mineral oil throughout the measurement. Imaging buffer was renewed every 12-15 hours.

### Data processing and image analysis

First, images were deconvolved by Huygens Professional version 18.04 (Scientific Volume Imaging, The Netherlands, http://svi.nl), using the CMLE algorithm (SNR:20, 40 iterations) run with a custom-made script written in Tcl/Tk. All further analysis steps were performed using a homemade analysis pipeline developed using MATLAB Release 2017b (The MathWorks, Inc., Natick, United States). First, we corrected for x-y drift in each cycle of hybridization. For each cycle *j*, the global x-y correction was obtained by cross-correlating the image of fiducial barcode *j* with that of the first barcode (reference cycle). This produced a single 3D vector for each barcode *j* and represented a ‘global’ correction applied to the whole FOV. Second, we used adaptive thresholding to pre-segment the spots of each fiducial barcode in each cell for all FOVs and for all barcodes. The 3D coordinates of each barcode were then found by using a 3D Gaussian fitting algorithm on the pre-segmented mask. Fiducial barcodes with sizes larger than the diffraction limit of light (∼2.2 pixels for our microscope) were then filtered out. Third, we obtained ‘local’ 3D correction vectors for each cell in each FOV. This was achieved by first using the global x-y correction vector to pre-align fiducial barcode spots in cycle *j* to fiducial barcode spots in the reference cycle. Then, we used image-based cross-correlation of these pre-aligned fiducial barcode images to reach sub-pixel accuracy in the correction vector. This approach allowed for 3D, subpixel accuracy drift-correction across the whole FOV. Forth, barcodes were segmented for all hybridization cycles in batch processing mode using optimized adaptive thresholding. 3D coordinated of each barcode were then determined by 3D Gaussian fitting of the segmented regions. These positions were corrected for drift by using the closest fiducial barcode vector obtained from the previous analysis step (local drift 3D correction, see above). Nuclei were segmented from DAPI images by adaptive local thresholding and watershed filtering. RNA images were segmented by manually drawing polygons over the nuclei displaying a pattern of active transcription. This was used to assign an expression status for each DAPI-segmented cell. Then, barcodes were attributed to each cell by using the DAPI segmentation. The efficiency of labeling per cycle for all barcodes was 60-70 percent (Figure S1F). The barcode localizations for each cell were then clustered as follows. First, the mean number of localizations per readout code was found and used as a measure of the maximum number of clusters N (1 cluster for paired chromosomes, 2 clusters for unpaired chromosomes, etc). Then, K-means was used to separate barcodes positions into N clusters. Finally, pairwise distances and contact frequencies were calculated for each cluster in each cell.

All image processing was carried out on Linux terminals connected to a server running Linux Ubuntu 16.04 Xenial or CentOS 7, with 32 CPU processors, two GeForce GTX 1080Ti GPU cards, and 128GB of RAM.

### Multiscale correlation of Hi-M distance maps

To compare Hi-M matrices at different scales, we did the following steps. First, we convolved maps with a Gaussian kernel of size s (standard deviation). Then, we calculated the correlation between maps and repeated this process for different values of s, ranging from 0.1 to 5. These values correspond to approximate genomic sizes ranging from 5 to 150 kb. As a control, we simulated random matrices with the same sizes and intensity ranges as the experimental matrices and repeated the same procedure. This process was repeated for 200 random matrices and the average correlation curves are shown in Figure 2B.

### Precision of the method

The mean localization accuracy after drift correction was 43 ± 21 nm in xy and 51 ± 58 nm in z (Figure S1E). This was obtained by measuring the distance between fiducial barcodes after applying the correction obtained from the previously described image correlation method. To further verify the precision of co-localization, a single locus was simultaneously labeled with encoding probes bearing binding sites for two distinct readout probes. After image registration, two sequential hybridization cycles with readout probes targeting the selected loci were performed. The co-localization precision after drift correction was of 83 ± 60 nm in xyz (Figure S1F). To further challenge the quality of drift correction and co-localization, a similar control experiment was performed but separating the readout hybridization cycles by 10 additional cycles and by letting the sample mounted in the microscope rest more than 24 hs. In these conditions, drift correction accuracy and precision of colocalization were equivalent to the previous control, ensuring that our internal marquer drift correction was unaffected by the set-up or sample stability during long acquisitions. As previously described (Cattoni et al., 2017), we calculate the absolute contact probability by integrating the area of the pairwise distance distribution below 120 nm. This threshold value was chosen from the integration of the pairwise distance distribution of a doubly-labeled locus.

### Hi-C and Chip-Seq data processing

Raw Hi-C sequencing data were processed using the scHiC2 pipeline (Nagano et al., 2017). Construction of expected models and Hi-C contact scoring was performed using the ‘shaman’ R package (https://bitbucket.org/tanaylab/shaman, (Cohen et al., 2017). Raw ChIPseq sequencing data were mapped to the dm3 reference genome using the bowtie.2 algorithm. Linear read density profiles at 10bp resolution where produced using MACS 1.4 (Zhang et al., 2008) after merging replicates. RNAseq RPM (reads per million) profiles where produced by aligning raw, paired-end, sequencing reads to the dm3 reference genome (BDGP R5/dm3 UCSC gene annotation) using STAR (Dobin et al, 2013) with ‘unstranded’ output.

We interpolated Hi-C matrices to make comparisons with Hi-M maps. For this, we extracted the contact frequencies at the genomic positions at which barcodes were located to construct an interpolated Hi-C map. Interpolated and full Hi-C maps displayed very similar features (Figures S1I, S3A).

## QUANTIFICATION AND STATISTICAL ANALYSIS

The number of nuclei quantified per experiment (N) are indicated in each of the corresponding figure legends.

## DATA AND SOFTWARE AVAILABILITY

Experimental datasets (lists of pairwise distances for nc 14, nc 9-13 mitosis, nc 12-13 interphase) and software package developed to analyze 3D deconvolved images produced by our Hi-M microscope have been uploaded to Mendeley Data: *http://dx.doi.org/10.17632/5f5hd9yj3z.1#folder-26d1f8c0-fc58-4b87-8c4f-cd8fa294a555*.

## ADDITIONAL RESOURCES

Table S4. Full Oligopaint library sequences, Related to STAR Methods.

## Supplementary figure legends

**Figure S1.**
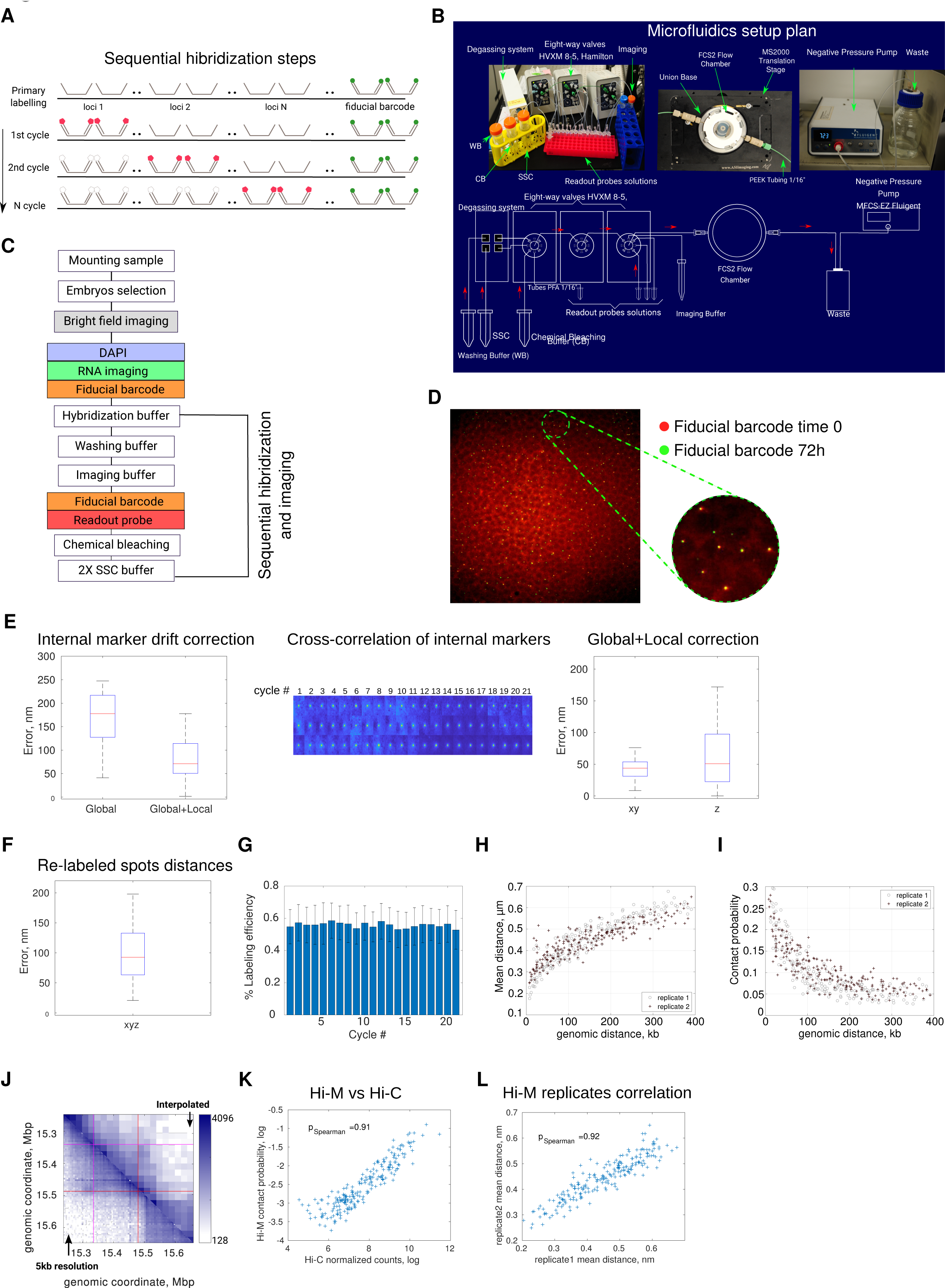
Details of the setup, the sequential hybridization and validation parameters of Hi-M, Related to Figure 1. A. Schematic representation of barcodes and sequential hybridization steps. The primary DNA hybridization using an oligopaint library was made at the bench. Next, hybridization with the fiducial barcode readout (insensitive to chemical bleaching) was performed also at the bench and was followed by a crosslinking step. Embryos were then immobilized within a microfluidics device. Each sequential hybridization cycle comprised the injection of a readout probe, incubation, imaging and chemical cleavage of the fluorophore from the readout probe. B. Schematic representation of microfluidics setup (lower panel) and images of the employed set-up (upper panel). The microfluidics setup was build according to (Chen et al., 2015) with modifications. C. Schematic representation of Hi-M acquisition pipeline. A first image registration phase comprised the imaging of DAPI-stained nuclei, of RNA probes, and of fiducial barcodes. Sequential hybridization/imaging cycles were performed within the robotic microscope and comprised the steps described schematically in the figure. D. Embryo stability at the beginning and end of a 72 h Hi-M acquisition. Embryos were attached to a poly-L-lysine-coated coverslip and mounted onto the microfluidics chamber. A representative field-of-view displaying a region of an embryo in the fiducial barcode channel is shown. Colors represent fluorescence intensity signal of fiducial barcode at 0h (red) and 72 h (green). Images were drift corrected using the protocol described in Methods. Inset displays a magnification of the selected area. E. <u>Left panel</u>, boxplot of the residual error in xyz between fiducial barcodes after the global and local drift correction. The global correction was obtained by cross-correlation of fiducial barcode images at different cycles. An improved local correction is obtained by: (1) segmenting fiducial barcodes in the first cycle; (2) subvolumes of size 20 × 20×60 pixels centered at each segmented object are extracted for the first cycle; (3) subvolumes are extracted for other cycles. Here the center of the subvolume is interpolated by using the localization of the fiducial mark in the first cycle and the global correction for each cycle obtained by image cross-correlation; (4) each subvolume *i* in cycle *j* is cross-correlated with subvolume *i* in cycle 1 to obtain a local correction vector specific for fiducial barcode *i*. A final correction vector for each segmented fiducial barcode and for each cycle is obtained by adding the global and local correction vectors. <u>Middle panel</u>: Examples of subvolumes for 3 fiducial barcodes (rows) are shown for all cycles. A typical field-of-view contains hundreds of fiducial barcodes. <u>Right panel</u>, boxplot of the residual error after the global and local correction in xy and z directions. Median values: 42 nm (xy) and 51 nm (z). F. To validate the drift correction method, we sequentially labeled and imaged a single barcode using two readout probes. Then, we calculated the distances the residual errors after global and local drift corrections and plotted them as a boxplot. The remaining mean xyz residual error was smaller than 90 nm. G. Mean labeling efficiencies for each imaging cycle. Labeling efficiency is defined as the number of nuclei having at least one detected barcode foci in the corresponding cycle over the total number of nuclei. Error bars correspond to the standard deviations of labeling efficiencies between different embryos. H-I. Mean physical distance (G) and absolute contact probability (H) vs. genomic distance from Hi-M for two replicates (different embryos at equivalent developmental timings, according to cell membrane invagination occuring during nc 14). Gray circles represent replicate 1 whereas brown crosses represent replicate 2. J. Replicate 1 mean pairwise distances vs replicate 2 mean pairwise distances. Replicate 1 and 2 are data from different embryos two at equivalent developmental timings. Pearson correlation coefficient = 0.92. K. Comparison between interpolated (upper triangle) and full resolution (5kb) Hi-C normalized maps (bottom triangle) for nc 14 embryos. Correlation between Hi-M contact probability and Hi-C normalized counts. Pearson correlation coefficient = 0.91.

**Figure S2.**
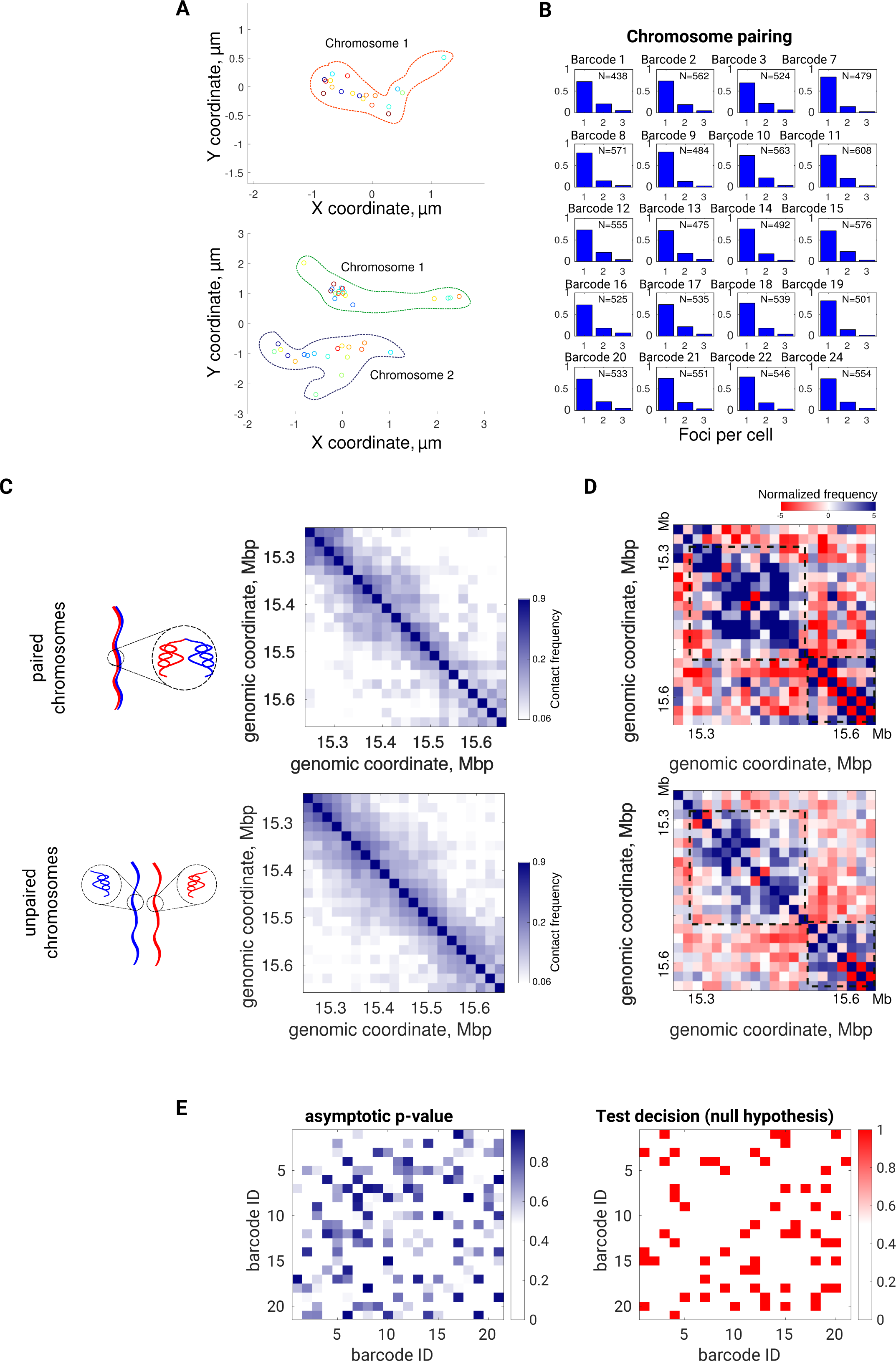
Paired and unpaired homologous chromosomes structure revealed by Hi-M, Related to Figure 2. A. Top view of an example of paired (top panel) and unpaired (bottom panel) chromosomes defined as in the main text. Circles represents the localization of the different barcodes in the XY space whereas dotted lines indicates the region supposedly occupied for each chromosome. B. Distribution in the number of spots detected per barcode in a representative nc 14 embryo. Each subplot represents a different barcode. Pairing was similar for all barcodes (>70%), consistent with previous studies (Cattoni et al., 2017; Fung et al., 1998). For representation purposes, only nuclei with up to 3 spots/barcode/nucleus are shown. The nuclei with four or more spots/barcode/nucleus was <3%. C. <u>Right panels</u>: Hi-M absolute contact probability maps for paired and unpaired chromosomes. <u>Left panels</u>: schemes of paired and unpaired chromosomes. D. Hi-M normalized contact probability maps of paired (upper panel) and unpaired (lower panel) chromosomes. Dashed black boxes represent TADs detected by Hi-C. E. KS test decision (left panel) and asymptotic p-value (right panel) maps obtained by comparing the pairwise distance distributions for each pair of barcodes for paired or unpaired chromosomes. Significance level p= 0.05. The results indicate that distance distributions between ‘paired’ and ‘unpaired’ conformations are different (i.e. do not pass the null hypothesis test) for most - but not all - pairwise barcode combinations. As with our correlation analysis between normalized mean distance matrices, we can conclude that the conformations are partially different -most notably in the region surrounding TADs.

**Figure S3.**
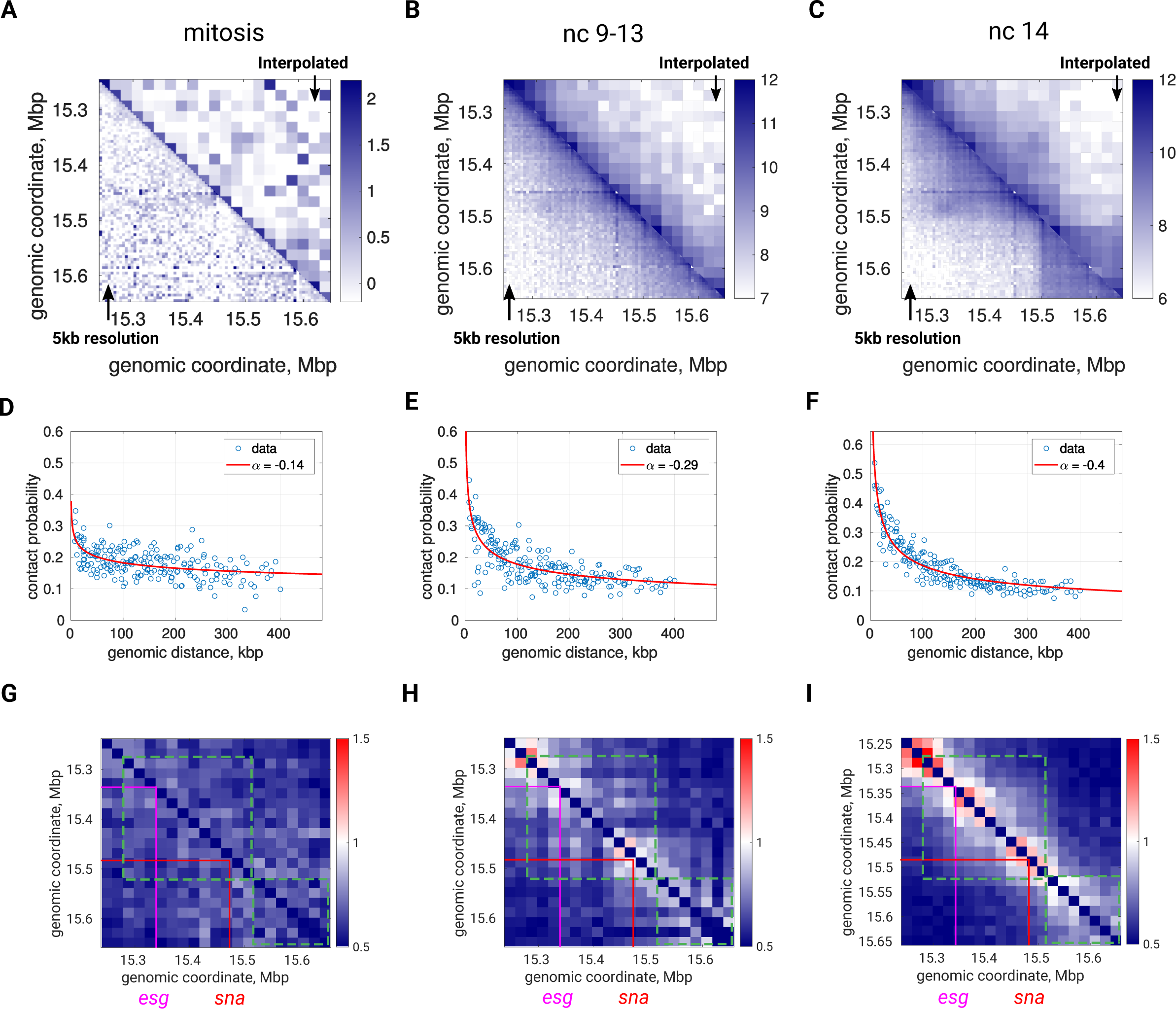
Changes in absolute contact probability and in the coefficient of variation during *Drosophila* development, Related to Figure 3. A-C. Comparison between interpolated (upper triangles) and full resolution (5 kb) Hi-C normalized maps (bottom triangles) for embryos in mitosis (A) nc 9-13 (B) and nc 14 (C). N= 1430 (mitosis), N=2933 (nc 12-13), N=6643 (nc 14). D-F. H-M absolute contact probabilities *vs*. genomic distances for embryos in mitosis (D), nc 9-13 (E) and nc 14 (F). Solid red lines are power-law fits with the scaling exponent indicated in the insets. G-I. Ratio between standard deviation and mean pairwise distance maps from embryos undergoing mitosis (G), nc 9-13 (H) and nc 14 (I). Scale bar on the right goes from 0.5 (blue), 0 (white) to 1.5 (red). N as in panels A-C.

**Figure S4.**
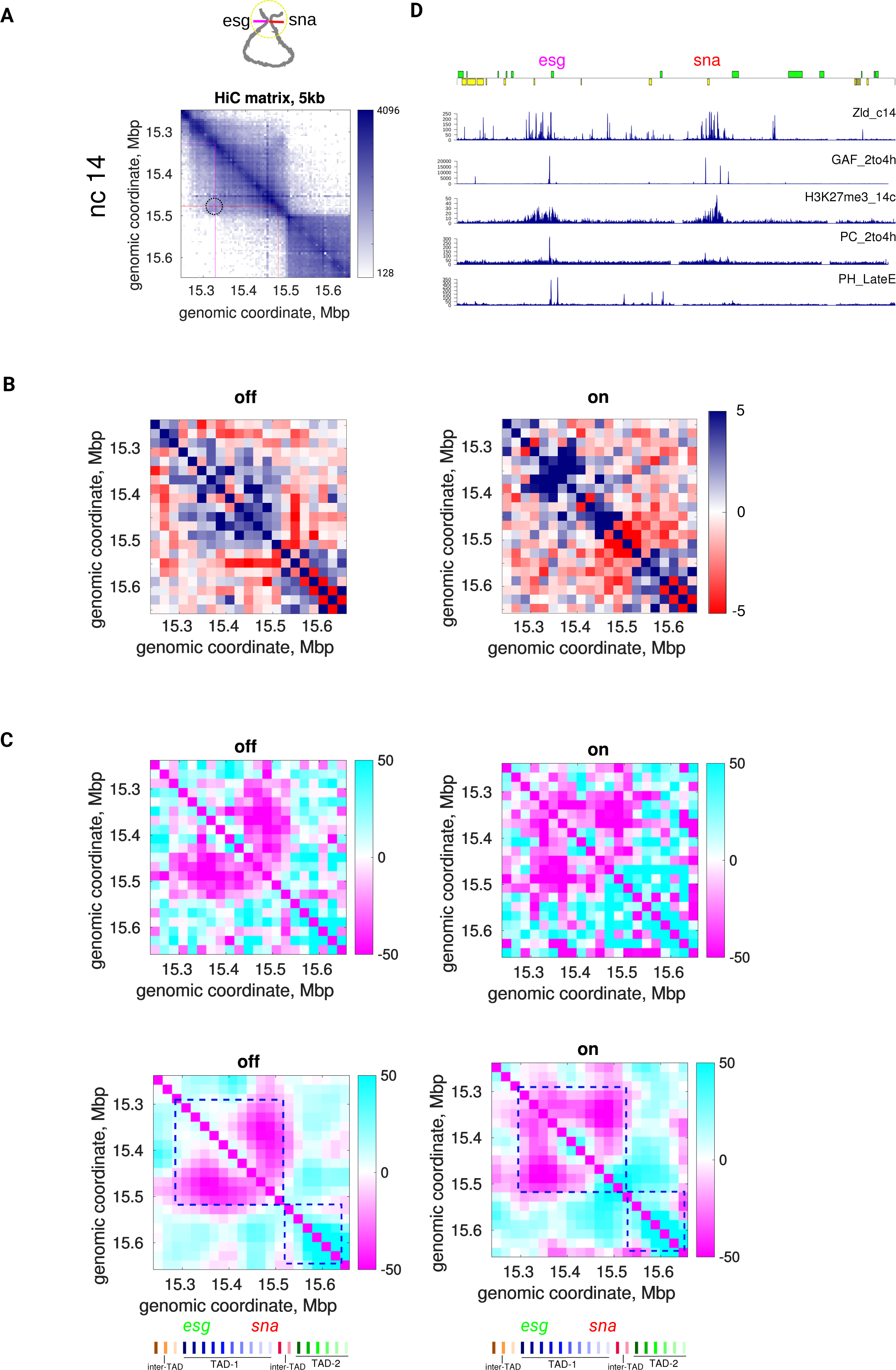
Correlation between changes in chromosome organization with transcription observed by Hi-M, Related to Figure 4. A. Full resolution (5kb) Hi-C map for nc 14 embryos. Relative contact frequency is color-coded according to scale bar (log scale). Black dotted circle indicates the position of the *sna-esg* loop. The positions of *esg* and *sna* are indicated with magenta and red lines, respectively. Top panel: schematic representation of the loop between *esg* and *sna*. B. Normalized Hi-M contact maps for nuclei in nc 14 for *sna* inactive (left panels) and *sna* active nuclei (right panels). Scale bar on the right indicates. C. Normalized Hi-M distance maps for nuclei in nc 14 for *sna* inactive (left panels) and *sna* active nuclei (right panels). Raw maps are shown in the upper panels and gaussian filtered panels are displayed in the lower panels. Barcodes are shown below. D. Right panel displays: genes, Chip-Seq data for Zelda in nc 14(Harrison et al., 2011), Chip-Seq for GAGA-factor (GAF) in stage 4-8 (Koenecke et al., 2016), Chip-Seq of H3K27me3 in nc 14 (Li et al., 2014), Chip-seq for polycomb and Ph (Polycomb group proteins) in nc 14 and stage 12, respectively (Koenecke et al., 2016; Schuettengruber et al., 2014).

