## Supplementary material for "Microscopy-based chromosome conformation capture enables simultaneous visualization of genome organization and transcription in intact organisms": Table

Table S1: Barcodes coordinates, Related to STAR Methods section

| Library | Oligopaint library | Universal Primers | Read out | Start coordinate | End coordinate | Coverage (kb) | # of Oligos | Rows # in Table 5 |
|---|---|---|---|---|---|---|---|---|
| 1 | 2L_42_oligo | BB287-FWD / BB288-REV | RT1 | 15244500 | 15251200 | 6.7 | 75 | 1-75 |
| 2 | 2L_42_oligo | BB287-FWD / BB288-REV | RT2 | 15269900 | 15275500 | 5.6 | 75 | 76-150 |
| 3 | 2L_42_oligo | BB287-FWD / BB288-REV | RT3 | 15331200 | 15337700 | 6.5 | 75 | 151-225 |
| 4 | 2L_42_oligo | BB287-FWD / BB288-REV | RT22 | 15370000 | 15376400 | 6.4 | 75 | 226-300 |
| 5 | 2L_42_oligo | BB287-FWD / BB288-REV | RT23 | 15421000 | 15427000 | 6 | 75 | 301-375 |
| 6 | 2L_42_oligo | BB287-FWD / BB288-REV | RT24 | 15433900 | 15440000 | 6.1 | 75 | 376-450 |
| 7 | 2L_42_oligo | BB287-FWD / BB288-REV | RT7 | 15454200 | 15460000 | 5.8 | 75 | 451-525 |
| 8 | 2L_42_oligo | BB287-FWD / BB288-REV | RT8 | 15474000 | 15480300 | 6.3 | 75 | 526-600 |
| 9 | 2L_42_oligo | BB287-FWD / BB288-REV | RT9 | 15499000 | 15504500 | 5.5 | 75 | 601-675 |
| 10 | 2L_42_oligo | BB287-FWD / BB288-REV | RT10 | 15548000 | 15554800 | 6.8 | 75 | 676-750 |
| 11 | 2L_42_oligo | BB287-FWD / BB288-REV | RT11 | 15598000 | 15604300 | 6.3 | 75 | 751-825 |
| 12 | 2L_42_oligo | BB287-FWD / BB288-REV | RT12 | 15644000 | 15652000 | 8 | 75 | 826-900 |
| 13 | 2L_42_oligo | BB287-FWD / BB288-REV | RT13 | 15255000 | 15259000 | 4 | 60 | 901-960 |
| 14 | 2L_42_oligo | BB287-FWD / BB288-REV | RT14 | 15263000 | 15267000 | 4 | 56 | 961-1016 |
| 15 | 2L_42_oligo | BB287-FWD / BB288-REV | RT15 | 15295000 | 15300500 | 5.5 | 75 | 1017-1091 |
| 16 | 2L_42_oligo | BB287-FWD / BB288-REV | RT16 | 15320000 | 15325500 | 5.5 | 75 | 1092-1166 |
| 17 | 2L_42_oligo | BB287-FWD / BB288-REV | RT17 | 15352000 | 15358000 | 6 | 60 | 1167-1226 |
| 18 | 2L_42_oligo | BB287-FWD / BB288-REV | RT18 | 15395000 | 15401300 | 6.3 | 75 | 1227-1301 |
| 19 | 2L_42_oligo | BB287-FWD / BB288-REV | RT19 | 15486500 | 15491500 | 5 | 59 | 1302-1360 |
| 20 | 2L_42_oligo | BB287-FWD / BB288-REV | RT20 | 15523000 | 15529000 | 6 | 75 | 1361-1435 |
| 21 | 2L_42_oligo | BB287-FWD / BB288-REV | RT21 | 15574000 | 15580300 | 6.3 | 75 | 1436-1510 |
| 22 | 2L_42_oligo | BB287-FWD / BB288-REV | RT25 | 15627000 | 15633000 | 6 | 75 | 1511-1585 |

Table S2: Readout region sequences, Related to STAR Methods section

| Readout | Sequence |
|---|---|
| RT1 | CACACGCTCTTCCGTTCTATGCGACGTCGGTG |
| RT2 | GACCAAGAGCGGACGTTGTGCCCAATGATCGC |
| RT3 | GACACGTGATCCGCGATACGATGAAAGCGCGA |
| RT7 | CGCAACGCTTGGGACGGTTCCAATCGGATC |
| RT8 | CGAATGCTCTGGCCTCGAACGAACGATAGC |
| RT9 | ACAAATCCGACCAGATCGGACGATCATGGG |
| RT10 | CAAGTATGCAGCGCGATTGACCGTCTCGTT |
| RT11 | AAGTCGTACGCCGATGCGCAGCAATTCACT |
| RT12 | CGAAACATCGGCCACGGTCCCGTTGAACTT |
| RT13 | ACGAATCCACCGTCCAGCGCGTCAAACAGA |
| RT14 | CGCGAAATCCCCGTAACGAGCGTCCCTTGC |
| RT15 | GCATGAGTTGCCTGGCGTTGCGACGACTAA |
| RT16 | CCGTCGTCTCCGGTCCACCGTTGCGCTTAC |
| RT17 | GGCCAATGGCCCAGGTCCGTCACGCAATTT |
| RT18 | TTGATCGAATCGGAGCGTAGCGGAATCTGC |
| RT19 | CGCGCGGATCCGCTTGTCGGGAACGGATAC |
| RT20 | GCCTCGATTACGACGGATGTAATTCGGCCG |
| RT21 | ACACCCTTGCACGTCGTGGACCTCCTGCGCTA |
| RT22 | AGAACGATCCAGCGAGATCAAGTGGAGCTGCG |
| RT23 | CATTGCCGTATGGGCTAGGATGACCTGGCTCG |
| RT24 | GCATTCACCCTTGCACGATACCGAGCCACACC |
| RT25 | GCCCGTATTCCCGCTTGCGAGTAGGGCAAT |

Table S3: Fluorescent Readout probes, Related to STAR Methods section

| Readout | Sequence |
|---|---|
| revRT1 | CACCGACGTCGCATAGAACGGAAGAGCGTGTG/iThioMC6-D//3AlexF647N/ |
| revRT2 | GCGATCATTGGGCACAACGTCCGCTCTTGGTC/iThioMC6-D//3AlexF647N/ |
| revRT3 | TCGCGCTTTCATCGTATCGCGGATCACGTGTC/iThioMC6-D//3AlexF647N/ |
| revRT7 | GATCCGATTGGAACCGTCCCAAGCGTTGCG/iThioMC6-D//3AlexF647N/ |
| revRT8 | GCTATCGTTCGTTCGAGGCCAGAGCATTCG/iThioMC6-D//3AlexF647N/ |
| revRT9 | CCCATGATCGTCCGATCTGGTCGGATTTGT/iThioMC6-D//3AlexF647N/ |
| revRT10 | AACGAGACGGTCAATCGCGCTGCATACTTG/iThioMC6-D//3AlexF647N/ |
| revRT11 | AGTGAATTGCTGCGCATCGGCGTACGACTT/iThioMC6-D//3AlexF647N/ |
| revRT12 | AAGTTCAACGGGACCGTGGCCGATGTTTCG/iThioMC6-D//3AlexF647N/ |
| revRT13 | TCTGTTTGACGCGCTGGACGGTGGATTCGT/iThioMC6-D//3AlexF647N/ |
| revRT14 | GCAAGGGACGCTCGTTACGGGGATTTCGCG/iThioMC6-D//3AlexF647N/ |
| revRT15 | TTAGTCGTCGCAACGCCAGGCAACTCATGC/iThioMC6-D//3AlexF647N/ |
| revRT16 | GTAAGCGCAACGGTGGACCGGAGACGACGG/iThioMC6-D//3AlexF647N/ |
| revRT17 | AAATTGCGTGACGGACCTGGGCCATTGGCC/iThioMC6-D//3AlexF647N/ |
| revRT18 | GCAGATTCCGCTACGCTCCGATTCGATCAA/iThioMC6-D//3AlexF647N/ |
| revRT19 | GTATCCGTTCCCGACAAGCGGATCCGCGCG/iThioMC6-D//3AlexF647N/ |
| revRT20 | CGGCCGAATTACATCCGTCGTAATCGAGGC/iThioMC6-D//3AlexF647N/ |
| revRT21 | TAGCGCAGGAGGTCCACGACGTGCAAGGGTGT/iThioMC6-D//3AlexF647N/ |
| revRT22 | CGCAGCTCCACTTGATCTCGCTGGATCGTTCT/iThioMC6-D//3AlexF647N/ |
| revRT23 | CGAGCCAGGTCATCCTAGCCCATACGGCAATG/iThioMC6-D//3AlexF647N/ |
| revRT24 | GGTGTGGCTCGGTATCGTGCAAGGGTGAATGC/iThioMC6-D//3AlexF647N/ |
| revRT25 | ATTGCCCTACTCGCAAGCGGGAATACGGGC/iThioMC6-D//3AlexF647N/ |
